# High-fidelity Musculoskeletal Modeling Reveals a Motor Planning Contribution to the Speed-Accuracy Tradeoff

**DOI:** 10.1101/804088

**Authors:** Mazen Al Borno, Saurabh Vyas, Krishna V. Shenoy, Scott L. Delp

## Abstract

The speed-accuracy tradeoff is a fundamental aspect of goal-directed motor behavior, empirically formalized by Fitts’ law, which relates movement duration to movement distance and target width. Here, we introduce a computational model of three-dimensional upper extremity movements that reproduces well-known features of reaching movements and is more biomechanically realistic than previous models. Critically, these features arise without the need of signal-dependent noise. We analyzed motor cortical neural activity from monkeys reaching to targets of different sizes. We found that the contribution of preparatory neural states to movement duration variability was greater for smaller targets than larger targets, and that movements to smaller targets exhibited less variability in preparatory neural states, but greater movement duration variability. Taken together, these results suggest that Fitts’ law emerges from greater task demands constraining the optimization landscape in a fashion that reduces the number of “good” control solutions (i.e., faster reaches). Thus, the speed-accuracy tradeoff could be a consequence of motor planning variability and optimal control theory, and not exclusively signal-dependent noise, as is currently held.

**Significance Statement:** A long-standing challenge in motor neuroscience is to understand the relationship between movement speed and accuracy, known as the speed-accuracy tradeoff. We introduce a computational model of reaching movements based on optimal control theory using a realistic model of musculoskeletal dynamics. The model synthesizes three-dimensional point-to-point reaching movements that reproduce kinematics features reported in motor control studies. Such high-fidelity modeling reveals that the speed-accuracy tradeoff as described by Fitts’ law emerges even without the presence of motor noise, which is commonly believed to underlie the speed-accuracy tradeoff. This suggests an alternative theory based on suboptimal control solutions. The crux of this theory is that some features of human movement are attributable to planning variability rather than execution noise.

## Introduction

Elite tennis players are particularly skilled at striking a balance between the speed and the accuracy of their serves. If they only focus on placing the ball in a strategic location, they may inadvertently advantage their opponent by not paying close enough attention to the speed of their serve. On the other hand, serving the ball with as much speed as possible is not desirable as the ball may fail to clear the net or land in a location that is advantageous to their opponent. This relationship, commonly known as the “speed-accuracy tradeoff,” is observed not only for purely motor, but also perceptual and cognitive tasks. In laboratory settings, the speed-accuracy tradeoff, quantified by Fitts’ law (Fitts 1954), is an empirical finding that movements that require greater accuracy tend to be slower than those that do not. Fitts’ law relates the width of the target (*W*), the movement distance (*A*) and the movement duration (*MD*): *MD* = *a* + *b*log_2_(2*A/W*), where *a* and *b* are subject-specific scalar parameters. The log_2_(2*A/W*) term is known as the “index of difficulty”.

A widely held theory, initially proposed by Harris and Wolpert, posits that signal-dependent noise (i.e., “motor noise” present at the neuromuscular junction) that scales with the size of the control signal is a sufficient theory to explain Fitts’ law (Harris & Wolpert 1998, Lunardini et al. 2015, McCrea et al. 2005, Peternel et al. 2018). The theory suggests that if motor noise increases with the size of the control signal, then moving rapidly, which requires large control signals, increases the variability in the final movement position. Conversely, reaching to a smaller target, with low variability in the final position, requires smaller control signals that consequently produce slower movements. Thus, this theory provides an explanation for the speed-accuracy tradeoff. In the present work, we show that the speed-accuracy tradeoff, as described by Fitts’ law, emerges even without the presence of motor noise.

A large body of prior work in this area has employed simple one-dimensional linear systems or torque-driven two-joint models of the arm that are restricted to moving in a plane (e.g., Flash et al. 1985, Fagg 2002, Harris 1998, Alexander 1997, DeWolf 2016, Sketch 2017). While it certainly is not the case that the mechanical properties of the plant being controlled explain phenomenon such as Fitts’ law, the patterns of neural activity required to drive goal-directed movements are necessarily constrained by the dynamics of the plant. When musculoskeletal dynamics are modeled with sufficient fidelity, and these musculoskeletal dynamics are controlled in an optimal fashion, motor behavior that is consistent with Fitts’ law emerges and thus provides a parsimonious theory of the speed-accuracy tradeoff.

Thus, a primary innovation in this work is the development of a computational model of upper extremity movements with a more realistic model of the musculoskeletal system than previously described (three-dimensional, 47 muscles and five degrees-of-freedom [Saul 2015]). The musculoskeletal model proposed by Saul and colleagues was previously used to track experimental kinematics, but not synthesize movements *de novo*. Our computational model synthesizes three-dimensional point-to-point reaching movements that reproduce features reported in motor control studies and in our experimental data. We focus on point-to-point movements because normal voluntary movements can be described as holding still after moving from one pose to another (Shadmehr 2017). The movements are synthesized by minimizing a cost function based on end-point accuracy and squared muscle activation terms. We use the squared muscle activation term as an approximation to metabolic power consumption (Winter 2009). We show that the asymmetry in the velocity profile with movement speed can be explained in terms of optimal control theory, as opposed to it either representing some unexplained feature, or resulting from constraints in the neural architecture (Bullock 1988), or a learned strategy when approaching a target (Beggs & Howarth 1972), or time-varying cost constraints (Nagasaki 1989), or signal-dependent noise with a minimum variance model (Tanaka et al. 2004).

Guided by this computational model, we propose a new theory: Fitts’ law emerges as a consequence of the difficulty associated with finding optimal control solutions for challenging tasks. Concretely, our theory results from the observation that during learning of a new task, the search space for finding the optimal control solution is very high-dimensional and contains many local minima. In our model, we use a stochastic optimizer, which attempts to find appropriate motor plans (i.e., reach trajectories) in this search space. Our choice of a stochastic optimizer is supported by the results of Churchland and colleagues that suggest that motor planning variability is a central source of movement variability (but also see Fig. 4E). As a consequence of the high dimensionality of the search space and the presence of local minima, the optimizer is more likely to find locally optimal solutions in lieu of globally optimal solutions. Our theory suggests that for more challenging tasks (e.g., reaching to smaller targets) the trajectory optimizer tends to find less effective, locally optimal solutions (i.e., slower reaches). For less challenging tasks (e.g., reaching to larger targets) the optimizer tends to find more effective, locally or globally optimal solutions (i.e., faster reaches). More challenging tasks constrain the search space: previously effective solutions are now discarded as they do not produce trajectories that reach the target.

A realistic musculoskeletal model allows us to study human movement variability with greater resolution. Our theory highlights the role of motor planning rather than execution noise to explain features of human movement. That is not to say that signal-dependent noise does not exist or could not play a part in Fitts’ law. Rather, given the results here, we must re-examine the assumption that signal-dependent noise is the key factor that gives rise to Fitts’ law. Reaching to a smaller target may exhibit smaller velocities to minimize the effects of signal-dependent noise. Alternatively, slower reach trajectories for smaller targets might be “easier” to find from an optimal control theory perspective. We believe our new theory provides a more complete account of the empirical observations, both behaviorally and neurally, surrounding the speed-accuracy tradeoff, without having to rely on assumptions surrounding improvements in the signal-to-noise ratio.

## Results

We first evaluated whether the computational model reproduces important kinematic features reported in motor control studies. In Fig. 1A-B, we compare the hand positions in a center-out fast reaching task from our model and the data reported in Beer et al. 2000. The simulations are synthesized with trajectory optimization (see *Trajectory Optimization*), specifying only the final target position and the movement duration, which we set to 350 ms. Note that our simulations reproduce the observation that arm reach trajectories (in humans and non-human primates) are gently curved, particularly near the targets. In Fig. 1C-D, we compare the velocity profile of our simulation with the data presented in Soechting 1984. We also compare the effect of the target size on the synthesized and experimental data in Fig. 2A-B. Soechting and colleagues report that a smaller target causes the movement to have an earlier peak velocity and a velocity that decays more rapidly. We reproduce both of these features in our simulations. The areas of the small and large targets are 2.55 cm^2^ and 3.54 cm^2^. The velocity profiles of our computational model are determined from an average of fifteen runs.

**Fig. 1.**
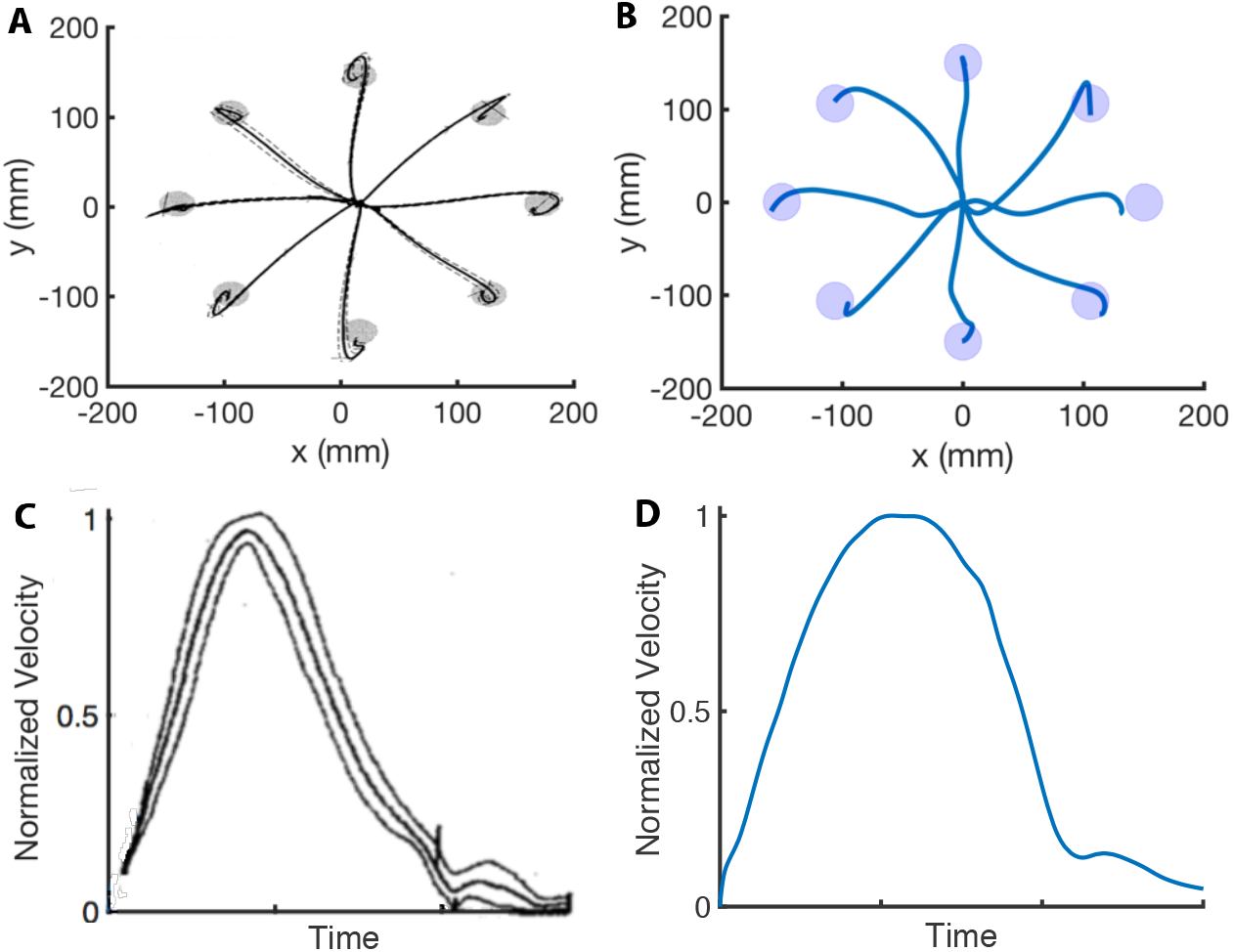
Center-out Reaching. **A-B:** Hand paths adapted from Beer et al. 2000 (**A**) compared to hand paths produced by our computational model (**B**) during a center-out fast reaching task with targets placed 150 mm away from the center position. **C-D:** Mean velocity profile (middle curve) in a reaching task adapted from Soechting 1984 (**C**) compared to the velocity profile produced by our computational model (**D**). We reproduce the bell-shaped velocity profile, including the smaller velocity peak at the end of the movement.

**Fig. 2.**
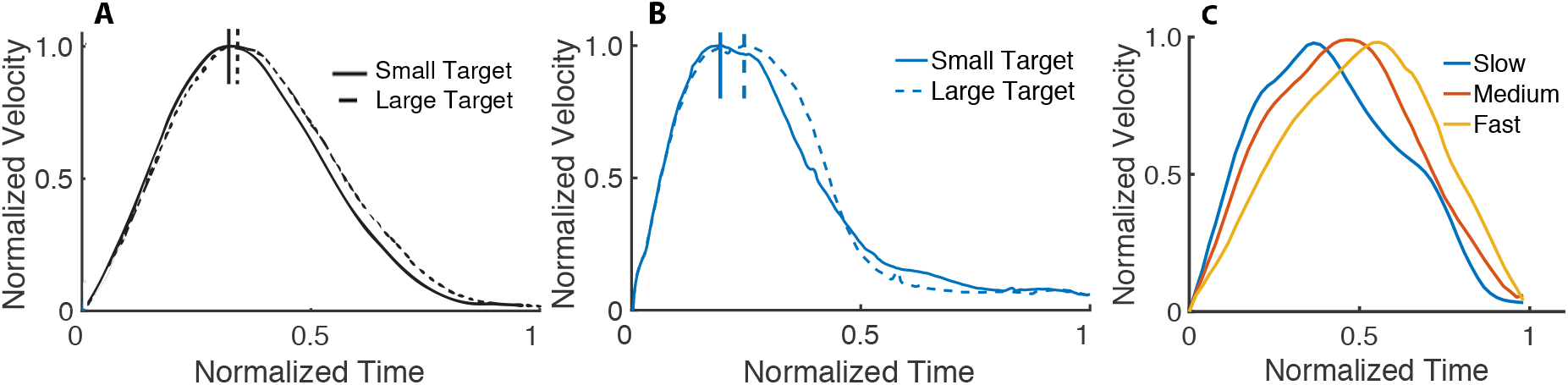
Velocity Profile based on Target Size and Speed. **A:** Effect of the target size in reaching movements. The dashed curve is the average velocity profile for the larger target, while the solid curve is for the smallest target (adapted from Soechting 1984). **B:** Average velocity profile of 15 runs from our simulations to a small and large target. All the curves are normalized to their peak velocity. **C:** Normalized velocity profiles (with normalized time) for fast, medium and slow movements. Our results are consistent with the observation (e.g., Nagasaki 1989) that slower movements tend to have an earlier peak velocity than faster movements.

Several studies have shown that the velocity profile is symmetric for intermediate speeds and becomes asymmetric with changes in speed (e.g., Nagasaki 1989, Ostry 1987). The ratio of the time to the peak velocity to the entire movement time tends to be smaller with slower movements. To study this, we optimized for twenty fast (0.18 s movement duration), medium (0.22 s movement duration) and slow (0.26 s movement duration) reaches to the same target. In Fig. 2C, we compare the normalized velocity profiles (with normalized time) between the best solutions (i.e., the trial with the lowest cost in Eq. (1) in *Materials and Methods*) for the movements at different speeds. Our model reproduces the speed-dependent asymmetry in the velocity profile. The relative peak velocities in Fig. 2C are 0.66, 0.51 and 0.4 for the fast, medium and slow movements, respectively. It has also been reported that the velocity profiles of slower movements tend to exhibit multiple local minima (e.g., Soechting 1984, van der Wel 2009); we replicate these findings in our simulations (see Fig. S2 in the Appendix for the velocity profile asymmetry for movement durations of 0.15 s, 0.25 s and 0.45 s). We performed the same experiment with a torque-driven model for movement durations of 0.2 s, 0.25 s and 0.3 s. In Fig. S3, we show that this simplified model does not reproduce the reported asymmetries in the velocity profiles.

In addition to the features reported in motor control studies, we compare the three-dimensional reaches performed by a typical subject and our computational model in Fig. S4 and in the accompanying video. Qualitatively, we observe that the movements resemble each other, with some discrepancies such as excessive or insufficient elbow flexion. In Fig. 3B, we compare the sagittal plane joint angles predicted by the model with our experimental data (see *Joint Kinematics Data*) when subjects are asked to reach a shoulder flexion 180° pose with the elbow extended. The RMSEs are 16.8° and 7° for the elbow and shoulder flexion angles.

**Fig. 3.**
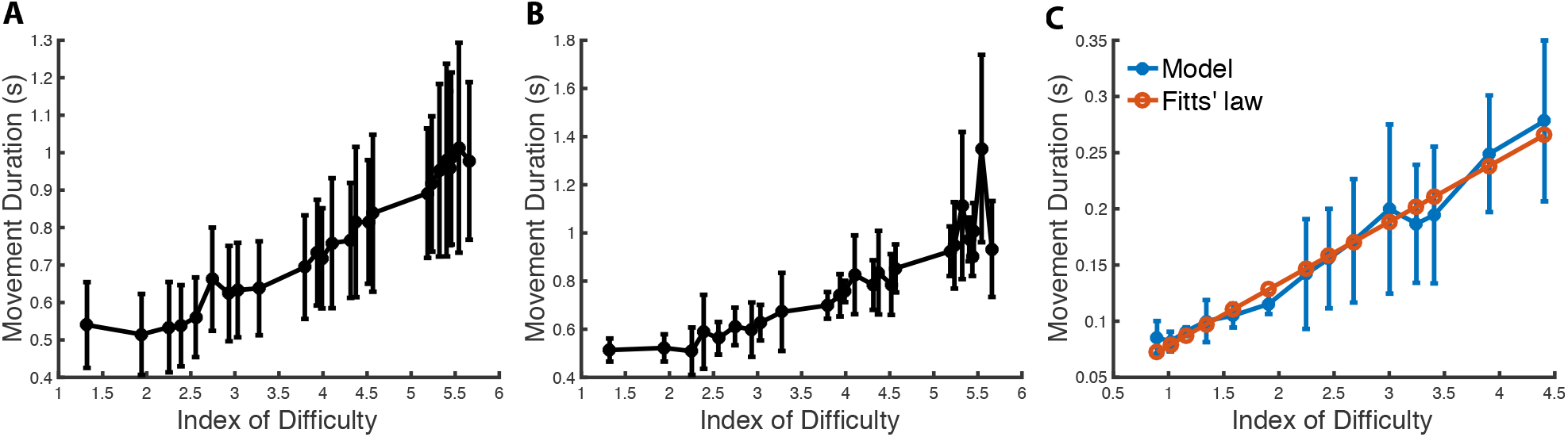
Speed-Accuracy Tradeoff. **A:** Movement duration depending on the index of difficulty from the dataset in Goldberg et al. (22). The mean and standard deviation are computed from 334 trials and 46 subjects (statistics computed across subjects). The experiment consists in subjects reaching with a mouse to targets of various distances and widths on a computer screen. **B:** Movement duration depending on the index of difficulty for a representative subject from Goldberg et al. (22) with statistics computed from seven trials to each target. **C:** Predicted movement duration to a target in a center-out reaching task when varying the target size. The x-axis is the width of the target in the direction of the movement, which varies from 0.16 m (index of difficulty: 0.9) to 0.014 m (index of difficulty: 4.41). The vertical bars are one standard deviation from the mean. We see that the model’s mean predictions are in close agreement with Fitts’ law (R^2^ = 0.974).

Next, we solved trajectory optimization problems to targets of different sizes to study the speed-accuracy tradeoff. We kept the distance fixed to **A** = 0.15 m and varied the target width from 0.16 m to 0.0141 m, which corresponds to an index of difficulty from 0.9 to 4.41. After determining the movement duration for different widths, we performed a least-squares fit to determine parameters *a* and *b* in Fitts’ law, obtaining values of 0.0234 and 0.0550, respectively. These values are within the range of values *a* ∈ [0.0047, 0.5239] and *b* ∈ [0.0393,0.1987] determined from the experimental data in Goldberg et al. 2015, where subjects reach with a mouse to targets of various distances and widths on a computer screen.

In Fig. 3C, we show that there is good agreement between the model’s predicted movement duration (averaged over ten runs) and Fitts’ law (R^2^ = 0.974). Note that the movement duration becomes more variable when the target size decreases as the optimization becomes more likely to fall in local minima. This feature is also present in the experimental data (see Fig. 3A-B), although it is not predicted by previous models (Harris & Wolpert 1998). Although the data in Goldberg et al. 2015 involves different experimental conditions than those of the simulated data of our center-out reach task, we expect some correlation when the index of difficulty is matched. This was verified as the correlation coefficient between our model’s mean predicted movement duration (as it varies with the index of difficulty) and the mean of the experimental data (Goldberg et al. 2015) is 0.959 (p < 0.001). The correlation coefficient between the standard deviations of our model and the experimental data is 0.70 (p = 0.0167). We performed the same experiment with a torque driven model (see Fig. S5). There is again good agreement between the model’s predicted mean movement duration (averaged over ten runs) and Fitts’ law (R^2^ = 0.88). However, the correlations between the predicted means and standard deviations and the experimental data in Goldberg et al. 2015 (see Fig. S5) are weak (R^2^ = 0.17 and R^2^ = 0.39, respectively) and not statistically significant.

We next evaluated the sensitivity of the results with respect to the trajectory optimizer. We repeated the experiments with a less effective trajectory optimizer by limiting the number of samples and iterations to 20 and 40 (see *Trajectory Optimization*), respectively. This translates to using about 12 times fewer samples during the optimization, so the computation time is about 12 times shorter. In Fig. S6, we see that model’s predicted movement duration means are still in close agreement with Fitts’ law (R^2^ = 0.969). The average means and standard deviations are 0.037 and 0.021, and larger than those in Fig. 3C. Our model does not include reaction time; therefore, as expected, it predicts faster movements than those that are observed experimentally. However, the relative movement durations (i.e., how the duration varies with the index of difficulty) in our model and the experimental data are still in close agreement. For example, the mean relative movement duration difference between an index of difficulty of 1.5 and 3.5 is about 1 s in both our model (see Fig. S5) and the experimental data (see Fig. 3A).

Given prior results regarding the potentially important role of neural preparatory activity during motor control (e.g., Afshar et al. 2011, Churchland et al. 2012, Ames et al. 2014, Shenoy et al., 2013), we examined the variability in the preparatory neural state during reaching movements to a small (0.75 cm) and a large target (2 cm). We measured neural activity from premotor and primary motor cortex of a Rhesus monkey (192 simultaneously recorded channels; 96 from each brain region), as he made reaching arm movements (Fig. 4A-B). Behaviorally, we found that reaches to smaller targets were slower than those to larger targets, as predicted by Fitts’ law (Fig. 4C). The movement durations to smaller targets were more variable than to larger targets (i.e., a standard deviation of 0.257 s compared to 0.234 s; p = 0.0262).

**Fig. 4.**
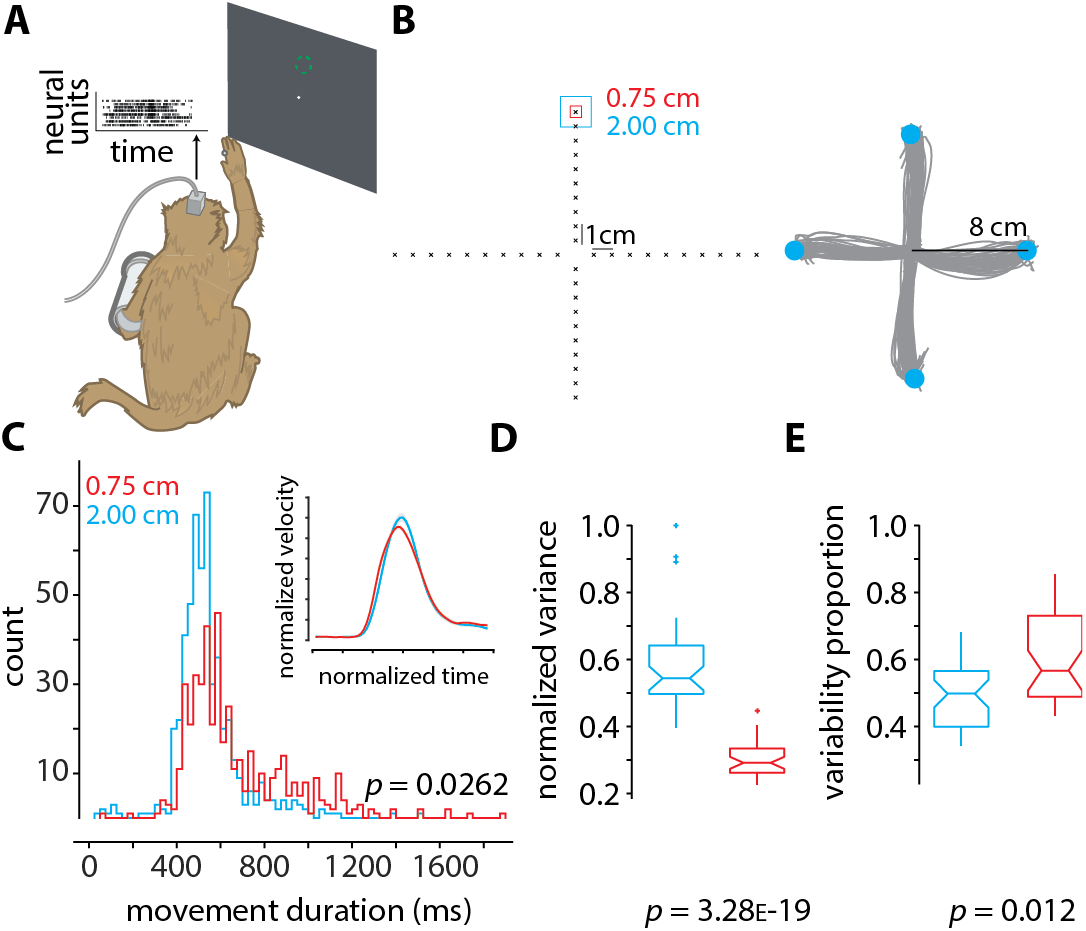
Neural Recordings and Analysis. **A:** Rhesus monkey made reaches in 3D space using his right arm, while neural activity from 192 channels was recorded from two Utah arrays surgically implanted into contralateral dorsal premotor and primary motor cortex. The monkey’s arm position in space controlled the velocity of the cursor on the screen (methods described in Even-Chen et al., 2019). **B:** (Left) the layout of the 40 targets to which the monkey made reaches. For each of the 40 targets, two conditions were presented in a block-wise format. In one condition (block) the target had an acceptance window of size 2 cm (cyan), and in the other the acceptance window was of size 0.75 cm (red). Within each block, the targets appeared randomly. Each block contained 400 trials (10 trials per target), and each experimental session contained about 10 blocks, 5 of each condition. (Right) representative arm reaches made by the monkey for four targets 8 cm away from the center with an acceptance window size of 2 cm. **C:** Histograms of movement duration for the large (2 cm) and small (0.75 cm) targets (inset shows the normalized velocity profiles). The observed difference in variance is statistically significant (p = 0.0262 via two-sample F-test). **D:** Normalized variance of the preparatory neural state for the small (0.75 cm targets, red) and large (2.00 cm targets, cyan) targets. The variance in the preparatory state is computed by first taking trial-averaged firing rates for the last 200 ms of preparation before the go-cue, performing principal components analysis, and finding the volume of the error ellipse in 3D space, which captures at least 80% of the total variance (similar to as described in Vyas et al., 2018 and Even-Chen et al., 2019). Note that we do not spike sort or assign spikes to individual neurons. We instead use threshold crossings (Trautmann et al., 2019). **E:** Proportion of movement variability explained by preparatory variability (as described in Churchland et al., 2006), for the small (0.75 cm targets, red) and large (2.00 cm targets, cyan) targets. In **D** and **E** the *p*-values are computed from one-way ANOVA.

Neurally, we found that reaches to smaller targets exhibited less variability in the preparatory state than reaches to larger targets (Fig. 4D). Intersecting this with findings that nearly half of movement variability can be attributed to preparatory variability for a highly practiced task (Churchland et al., 2006), we found that the contribution of preparatory variability to movement variability was larger for smaller targets than larger targets (Fig. 4E). Thus, these results establish an association (albeit correlative) between Fitts’ law and variability in neural activity during motor preparation.

## Discussion

In this study, we investigated a fundamental phenomenon underlying goal-directed motor control. Fitts’ law formalizes the notion that movement time is a function of both the distance to a target, and the size of that target. Taken together with a widely held theory initially proposed by Harris and Wolpert (Harris & Wolpert 1998), Fitts’ law is thought to arise from the fact that signal-dependent noise governs a person’s ability to move with greater accuracy towards smaller targets by reducing movement speed. Signal-dependent noise or “motor noise” has been shown across a variety of experiments, where noise scales in proportion to the size of the motor command being generated (e.g., Jones et al. 2002, Matthew et al. 1996). Thus, to a first-order approximation, signal-dependent noise is a sufficient theory to explain Fitts’ law. Here, we re-examine the assumption that Fitts’ law emerges primarily as a *consequence* of signal-dependent noise.

We developed a computational model of three-dimensional upper extremity movements that is more biomechanically realistic than previous work. Our model replicates kinematic features reported across several motor control studies. When solving the optimal control problem, we found that some of our simulations reproduced the empirical observation that arm movement paths are nearly straight and gently curved near the targets for fast reaches. The corresponding velocity profiles are bell-shaped and have a secondary, smaller peak near the target. These features are usually interpreted as discrete, open-loop corrections to acquire the target (Doeringer & Hogan 1998). While our results are consistent with this interpretation, our analyses also show that these features are present in some local minima when solving the optimal control problem. Out of 20 trials during a reaching task, we noted that the second best solution (measured with respect to the cost; Eq. 1) had these features, while the best solution consisted of a nearly straight trajectory. The curve and the secondary velocity peak at the end of the movement are a consequence of the hand reaching the target with a high velocity; the final, gentle curve, brings the hand velocity towards zero, minimizes “effort”, and keeps the hand inside the target.

Bullock and Grossberg 1998 have proposed that the speed-dependent asymmetry in the velocity profile arises due to neural dynamics and that models based on optimal control always predict symmetric velocity profiles. Our work demonstrates that asymmetric velocity profiles in reaching movements arise from optimal control methods with realistic biomechanical models, without relying on other modeling assumptions such as time-varying jerk constraints (Nagasaki 1989). This feature did not emerge in earlier work using simplified biomechanical models (Guigon 2007, Harris & Wolpert 1998). The upper extremity must decelerate when approaching the target to achieve zero velocity. The elbow flexors undergoing an eccentric contraction have a larger force generating capacity at higher speeds due to the force-velocity relationship of muscle (Winter 2009), which explains why faster movements have a later peak velocity.

During motor learning, there is a shift in performance that allows movements to be performed faster and more accurately. Previous studies hypothesize that this shift in performance is a result of improvements in signal-to-noise ratio (i.e., the ability to use large control signals without large motor noise). This is perhaps possible due to expanded neural representations and/or fine-tuning of individual neurons (Krakauer & Mazzoni 2011, Shmuelof et al. 2012). Indeed, there is evidence of an improved signal-to-noise ratio during motor skill learning in rodents (Kargo & Nitz 2004), but their mechanism proposes a reduction in signal-independent noise and pre-burst firing rates. If signal-dependent motor noise alone accounts for the speed-accuracy tradeoff, the mechanism that explains the shift in performance due to practice is not yet fully understood. Our analyses reveal that more stringent task constraints reduce the space of possible control solutions, thereby removing effective local optima from the search landscape (see Fig. 5). The controller (i.e., the central nervous system) is unable to find the global optimum solution for every task. Thus, for challenging tasks, the patterns of neural activity found during initial optimization (i.e., learning) correspond to local optima, leading to suboptimal behavior, e.g., reaching slower to smaller targets, consistent with Fitts’ law. Learning can force the motor system to explore different solutions, and if better local optima are found, the internal models are updated accordingly resulting in behavioral improvements (Haith & Krakauer 2018). Hence, our theory does not rely on assumptions regarding improvements in signal-to-noise ratio to explain the shift in performance with practice.

**Fig. 5.**
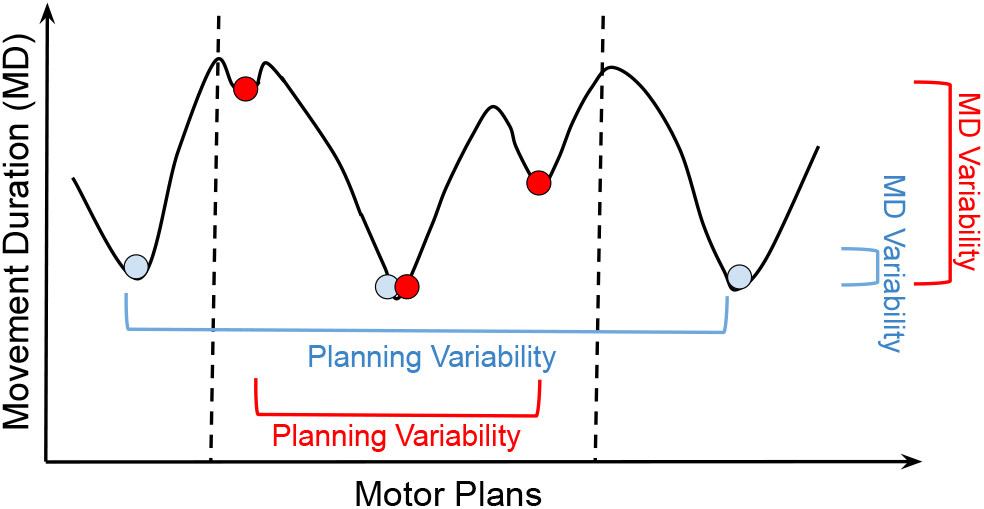
Optimization Landscape. Cartoon of how different motor plans can generate movements (i.e., driven by muscle excitation patterns) with different durations. The blue and red circles are examples of control solutions for the large and small targets, respectively. The dashed lines illustrate the constraints in the optimization landscape for the smaller target. From the analysis in Fig. 4C-D, for reaches to a small target (red), movement planning variability decreases (x-axis), but movement duration (MD) variability increases (y-axis). For reaches to a larger target (blue), movement planning variability increases, but movement duration variability decreases. According to our theory, reaches to larger targets are faster because of the presence of good local minima in the optimization landscape. We empirically observe that these good local minima have movement durations that are only slightly larger than the global minimum solution (i.e., the fastest reach to the target). A bad local minimum corresponds to a significantly slower reach. On average, reaches to the small target are slower because tighter constraints remove good local minima from the optimization landscape, which is otherwise identical to the landscape for the large target. While this decreases motor planning variability, it increases movement duration variability. In stochastic optimization, it is empirically found that having a large number of good local minima helps avoid bad local minima (Goodfellow et al. 2016, Chapter 8).

While our theory does not account for signal-dependent noise, the broader mechanism for motor control likely engages both the mechanism proposed here and certain aspects of signal-dependent noise. We posit that a realistic biomechanical model, as described here, allows for more accurate estimation of the optimization landscape (i.e., how the local optima are situated with respect to the control solutions). We have shown that a simplified torque-driven model does not produce results that are strongly correlated with prior experimental findings. Our experiments show that Fitts’ law emerges both with a standard and a less effective optimizer. This follows from the fact that for an optimizer with a given performance (i.e., measured by the number of samples and trials that it can explore), imposing progressively more stringent accuracy constraints removes good local optima from the optimization landscape, resulting in performance degradation and thus yielding Fitts’ law. A theory based purely on signal-dependent motor noise does not predict that, as seen in the experimental data, more severe accuracy constraints cause more movement duration variability. From this perspective, our theory provides a more complete account of the speed-accuracy tradeoff.

If our theory is correct, then the muscle excitations, activations and joint kinematics predicted for a suboptimal solution should be consistent with what is empirically observed in naïve or “non-expert” human participants. At the same time, our theory predicts that the globally optimal solution would result in muscle patterns that are consistent with empirical observations of highly-trained or “experts” performing the task. Future studies can perform this stringent test of our theory by utilizing our modeling framework to find the optimal solution for some subset of challenging tasks and compare them against muscle activity measured in a laboratory setting.

In addition to planning variability, dynamics also play a role in the speed-accuracy tradeoff. The differences in durations between the fastest trials to the smallest and largest targets provides an indicator of the role of dynamics in the speed-accuracy tradeoff. The fastest trial approaches the limit of what is dynamically feasible given enough trials. In our computational experiments, this difference amounts to 0.04 s. The average difference from the experimental data is also 0.04 ± 0.03 s (see Table S1). The remaining difference between the mean durations to the two targets (65% of the total difference in Fig. 3C) could be attributed to motor planning variability, which is the dominant factor behind the speed-accuracy tradeoff in our results. The proportions would likely be reversed if the target sizes were drastically different (e.g., comparing targets that were an order of magnitude apart in size). Our analysis does not preclude the possibility that signal-dependent noise is a factor in the speed-accuracy tradeoff in addition to dynamics and planning variability, but it shows that it is not a necessary condition and may not be the dominant factor as previously thought. The data in Table S1 also shows that the subjects that achieved the smallest difference in durations between the two targets on their best trials also had the largest variability (see subjects 1, 2 and 3). We hypothesize that this variability could be attributed to exploration, which allows for finding better control solutions, consistent with the experimental data. This is also consistent with recent findings that learning is associated with increases in task-relevant variability (Wu et al., 2014).

Turning our attention to neural mechanisms underlying motor cortical function, a current theory suggests that movement period neural activity evolves from an initial condition (termed the neural population preparatory state) set in premotor and primary motor cortex (Shenoy et al., 2013). Different movements have different initial conditions, and generating initial conditions closer to the optimal initial condition results in behavioral benefits (Afshar et al. 2011). Furthermore, trial-by-trial behavior variability is strongly linked to neural variability during motor preparation, even for highly practiced movements (Churchland et al. 2006), and motor learning is strongly linked with systematic changes to neural activity during motor preparation (Vyas et al. 2018). If movement planning variability is a dominant factor underlying the speed-accuracy tradeoff, then the proportion of movement duration variability explained by preparatory variability should increase with reaches to smaller targets (otherwise motor planning does not contribute to the speed-accuracy tradeoff). We have experimentally verified this prediction (see Fig. 4E). Our experiments here further show that reaches to smaller targets exhibit slower reaches and less variability in preparatory activity than reaches to larger targets (see Fig. 4D), while the movement duration exhibits more variability in smaller targets than in larger targets. These results are consistent with our theory (see Fig. 5), which suggests that some of the effective control solutions (i.e., motor plans for fast reaches) have to be discarded when reaching to smaller targets (i.e., variability in preparatory activity decreases, while movement duration variability increases). We note, however, that this is an initial *theory*, and future studies will need to evaluate its range of applications under additional experimental conditions.

As our theory relies largely on the motor plan it is tempting to link variability in the stochastic trajectory optimizer to variability in preparatory activity, especially given our findings in Fig. 4, which links Fitts’ law to variability in preparatory activity. While the preparatory state does not encode the whole reach trajectory, it does encode important parameters of the upcoming reach, such as distance, direction, speed, etc. (Messier & Kalaska 2000, Even-Chen et al., 2019). The results here, and those by Churchland and colleagues (Churchland et al. 2006), point to preparatory activity as also being a central source of variability, rather than signal-dependent noise alone being the key factor. Taken together, one tantalizing interpretation of these results is that Fitts’ law is tightly linked to movement planning (which includes many of the aspects of the preparatory activity analyzed here) and not just movement execution. The difficulty associated with finding optimal solutions for challenging tasks could then be akin to finding the optimal preparatory neural states (as well as learning the appropriate movement period dynamics), where task constraints govern the optimization landscape, as suggested by our theory. This provides a concrete avenue for future studies to further investigate the relative contribution of signal-dependent noise and motor planning in giving rise to the speed-accuracy tradeoff.

## Author contributions

MAB designed the study, performed the computational modeling and analyses, and wrote the manuscript. SV performed the animal experiments and analyses, helped interpret results, and draft the manuscript. KVS supervised the animal experiments and analyses and provided edits on the manuscript. SLD supervised all aspects of this work and provided edits on the manuscript.

## Acknowledgments

We thank Adrian Haith for very helpful comments on the manuscript. We thank Mackenzie Risch and Michelle Wechsler for expert surgical assistance and veterinary care. We thank W. L. Gore Inc. for donating Preclude artificial dura used as part of the chronic electrode array implantation procedure. We thank Dr. Stephen I. Ryu for surgical assistance. S.V. was supported by an NIH F31 Ruth L. Kirschstein National Research Service Award 5F31NS103409-02, an NSF Graduate Research Fellowship, and a Ric Weiland Stanford Graduate Fellowship. K.V.S. was supported by the following awards: National Institutes of Health (NIH) National Institute of Neurological Disorders and Stroke (NINDS) Transformative Research Award R01NS076460, NIH National Institute of Mental Health Grant (NIMH) Transformative Research Award R01MH09964703, NIH Director’s Pioneer Award8DP1HD075623, Defense Advanced Research Projects Agency (DARPA) Biological Technology Office (BTO) “REPAIR” award N66001-10-C-2010, DARPABTO “NeuroFAST” award W911NF-14-2-0013, the Simons Foundation Collaboration on the Global Brain awards 325380 and 543045, ONR and the Howard Hughes Medical Institute. S. L. D. was supported by the Mobilize Center, a National Institutes of Health Big Data to Knowledge (BD2K) Center of Excellence through Grant U54EB020405.

## Materials and Methods

### Modeling and Simulation

Our computational model uses the upper extremity biomechanical model introduced by Saul et al. 2015 and available at https://simtk.org/projects/upexdyn. The model consists of 47 Hill-type muscle-tendon actuators (Zajac 1989) with parameters and paths derived from experimental and anatomical studies. The skeletal model has five degrees-of-freedom representing the shoulder (elevation plane, elevation angle and shoulder rotation), the elbow (flexion/extension), and the forearm (pronation/supination). We have locked the wrist and finger joints. The simulation was performed with OpenSim (Delp 2007) and the Simbody physics engine (Sherman 2011) with a semi-explicit Euler integrator (accuracy 1e−2).

### Trajectory Optimization

Numerical methods for solving trajectory optimization problems is an on-going research topic (e.g., Al Borno 2013, Posa 2013, Todorov 2005). Our work adapts the method of Al Borno et al. 2013 to optimize the movement of musculoskeletal systems. We optimize for a time-indexed cubic B-spline that provides the muscle excitations to produce movement. The optimization is performed with Covariance Matrix Adaption (CMA-ES; Hansen 2006). The free variables are the spline knots muscle excitations values at every 0.1 s interval in the movement. We use 500 iterations and a population size of 20 in CMA-ES, which takes about three hours of computation time (after parallelization) with two Intel Xeon CPU E5-4640 processors. We initialize the optimization with all the knots set to 0.1. Given excitations, we use a forward simulation to obtain a trajectory, which is evaluated with respect to the following cost function:

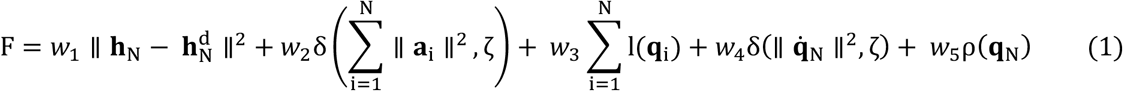

where **h**_N_ and 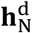, denote the actual and desired position of the center of the hand at the last timestep N of the trajectory, **a**_i_ denotes the vector of muscle activations at timestep *i*, l(**q**_i_) is a quadratic penalty on joint limit violations given pose **q**_i_, and 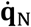 denotes the joint velocities at the last timestep, ρ(**q**_N_) is a quadratic term to encourage pronation of the forearm at the last timestep (as is typical in reaching movements to a target), and ∥ · ∥ denotes the 2-norm. It is challenging to tune the weights in the cost function as objectives tend to “fight” each other. For this reason, we introduce the δ(·, ·) function:

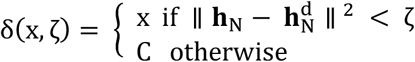

where we set the scalars ζ to 0.025 and C to 10^6^ (i.e., C ≫ x).This ensures that the optimization priorities finding a trajectory with the hand reaching close enough to the target (i.e., within 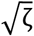 m) before considering other objectives, namely minimizing the sum of activations squared and achieving a final zero velocity. We tuned the weights *w*_1_, *w*_2_, *w*_3_, *w*_4_ and *w*_5_ to 5, 1e−3, 1e−5, 0.15 and 0.5. The cost function encourages the hand to be as close as possible to a desired target with a small velocity and the forearm in pronation, while minimizing muscle activations and joint limit violations throughout the movement. When the objective is to reach a target anywhere within a radius *r*, then we simply set *w*_1_ to zero when 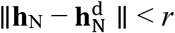.

The movement duration T is chosen based on experimental data or through optimization to study the speed-accuracy tradeoff. For the latter, we encourage the movement to reach the target as soon as possible by including the objective φ(**h**_i_) in the cost function (Eq. 1). We set φ(**h**_i_) to 0 if the target is reached at timestep *i*; otherwise, we set it to 1 (i.e., a penalty at each timestep where the hand position is outside the target). We perform the trajectory optimization with a fixed temporal horizon and the movement duration is determined when the hand position is within the target. Haith et al. 2012 argue that a hyperbolic cost of time objective term better accounts for the durations of saccades, but this remains a topic of investigation for other body movements. After setting the duration of the movement T for the integration, the number of timesteps N in the trajectory is chosen to achieve the desired accuracy in Simbody. For the experiments with our torque-driven model, we optimize for the torques, which we limit to ±50 Nm, instead of muscle excitations.

### Joint Kinematics Data

To validate our computational model, we collected three-dimensional upper extremity movement data. Right-handed subjects (n=4, age = 27 ± 3, mean ± std, 2 males, 2 females) gave written informed consent approved by the Stanford University Institutional Review Board. The subjects are asked to reach for desired final end poses (e.g., with shoulder abduction 90°, shoulder flexion 90° and 180°, etc.) from different initial positions. In Fig. S1, we show an example of an initial (A) and final position (B). Subjects are instructed to reach the target pose at a comfortable speed, without moving their trunk or feet. Upper extremity kinematics are measured with inertial motion capture (Roetenberg et al. 2009). Movement duration and the final hand position (computed from forward kinematics) are then used for motion synthesis with trajectory optimization, using the final hand position for the target position 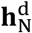.

### Fast Reaching Task

We collected experimental data from five right-handed subjects (n=5, age =28 ± 4.9, mean ± std, 4 males, 1 female) that gave informed consent approved by the Stanford University Institutional Review Board. The subjects performed reaches to a target, always starting from the same initial pose with the hand on the table. Subjects were instructed to reach the target as fast as possible. The duration of the reach is measured from an accelerometer attached on their hand (Xsens MTw Awinda with a 1000 Hz sampling frequency). Subjects performed ten reaches to a large square target (8 cm width) and to a small one (2 cm width). The movement amplitude was set to 15 cm. The data is presented in Table S1.

### Data Availability

The source code for the computer simulations and our data are available at https://simtk.org/projects/ue-reaching. Further information and requests for experimental monkey data pertaining to the results in this study should be directed to and will be fulfilled by Saurabh Vyas upon reasonable request.

