## Supplemental Material for "High-fidelity Musculoskeletal Modeling Reveals a Motor Planning Contribution to the Speed-Accuracy Tradeoff"

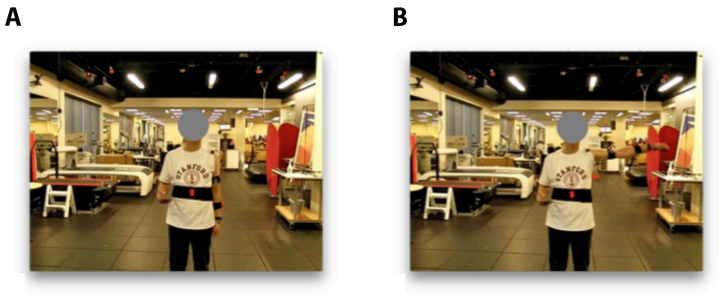


**Fig. S1. Data Collection. A-B:** Subjects are asked to move to a desired final pose (**B**) from a given initial pose (**A**). This figure is an example of a shoulder 0°-90° abduction movement.

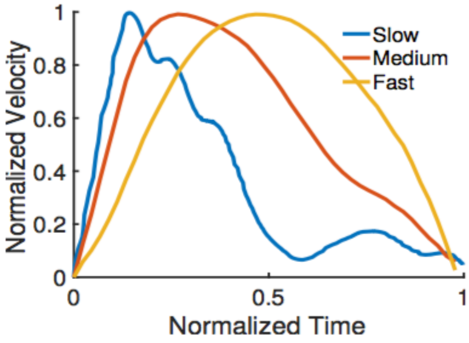


**Fig. S2. Asymmetry of the Velocity Profile based on Speed.** Normalized velocity profiles (with normalized time) for fast (0.15 s movement duration), medium (0.25 s movement duration) and slow movements (0.45 s movement duration). an earlier peak velocity than faster movements. Our model reproduces the speed-dependent asymmetry in the velocity profile. The relative peak velocities are 0.6133, 0.41 and 0.19 for the fast, medium and slow movements, respectively.

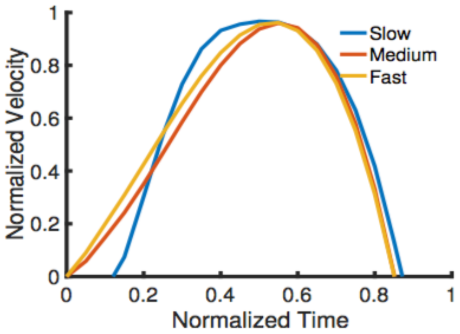


**Fig. S3. Asymmetry of the Velocity Profile based on Speed with a Torque Driven Model.** Normalized velocity profiles (with normalized time) for fast, medium and slow movements (20 s, 25 s, and 30 s movement durations, respectively). The torque driven model does not reproduce the observation that faster movements tend to have a later peak velocity than slower movements, while this was reproduced with our realistic muscle-based model (see Fig. 2C).

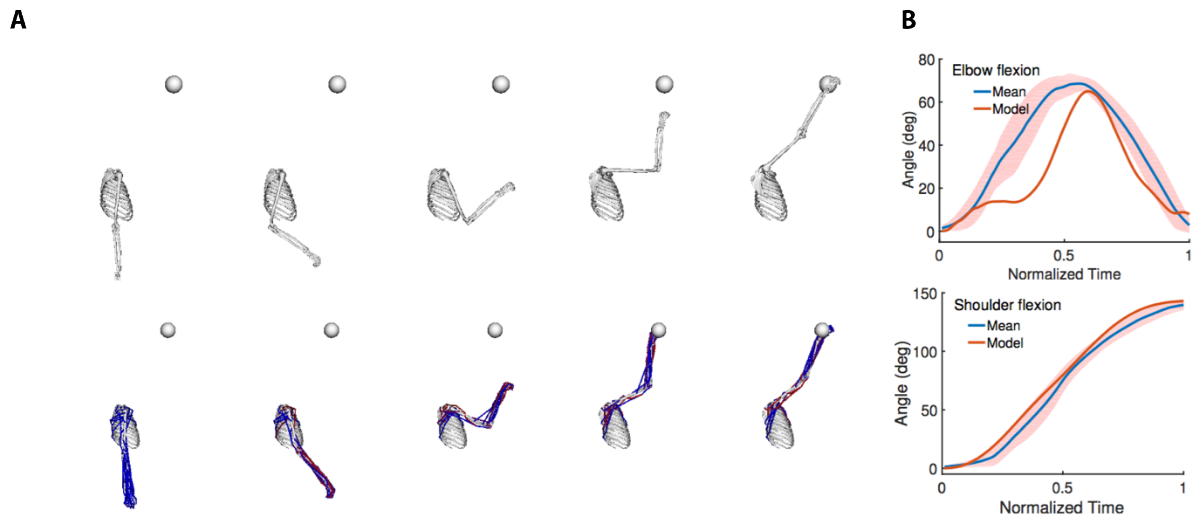


**Fig. S4. Movement Comparison. A:** We compare the reaches to a target (the sphere) of a representative subject (top; model without muscles) with the reaches synthesized by our computational model (bottom; model with muscles). **B: Joint Kinematics.** We plot the sagittal joint angles in the reaching movement. We compare the elbow and shoulder flexion angles over time from our model and the means from experiments. The shaded region is one standard deviation from the mean

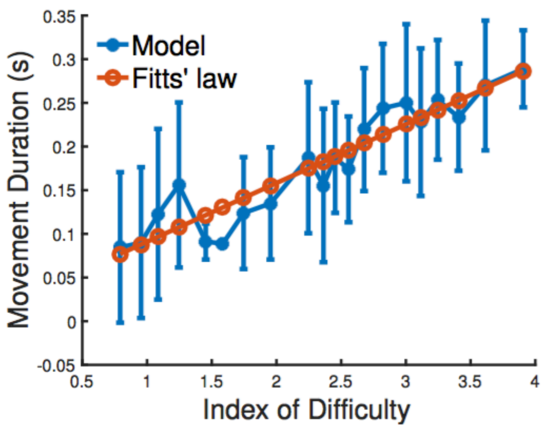


**Fig. S5. Speed-Accuracy Tradeoff with a Torque Driven Model.** Predicted movement duration to a target in a center-out reaching task when varying the target size with our torque driven model. The x-axis is the width of the target in the direction of the movement, which varies from 0.02 m (index of difficulty: 3.90) to 0.2 m (index of difficulty: 0.09). The vertical bars are one standard deviation from the mean. We see that the model's mean predictions are in close agreement with Fitts' law ($R^2 = 0.88$). Qualitatively, we see that unlike our muscle-based model (see Fig. 3C), the results with the torque driven model do not clearly predict the increase in movement duration variance with more severe task constraints. Both the predicted means and standard deviations are not statistically correlated with the experimental data in Fig. 3A.

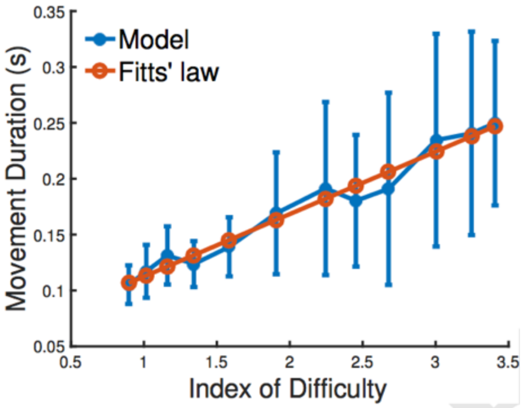


**Fig. S6. Impact of the Optimizer on the Speed-Accuracy Tradeoff.** We use a much less effective trajectory optimizer than in Fig. 3C to predict the movement duration to a target in a center-out reaching task when varying the target size. The x-axis is the index of difficulty of the reaching movement. We see that the model's mean predictions are still in very close agreement with Fitts' law ($R^2 = 0.969$).

| Subject | L mean(std) | L (min, max) | S mean(std) | S (min,max) |
|---|---|---|---|---|
| 1 | 0.19(0.03) | (0.16,0.24) | 0.29(0.13) | (0.18,0.59) |
| 2 | 0.25(0.06) | (0.11,0.31) | 0.27(0.11) | (0.12,0.38) |
| 3 | 0.18(0.04) | (0.10,0.24) | 0.25(0.06) | (0.11,0.31) |
| 4 | 0.26(0.04) | (0.19,0.34) | 0.32(0.03) | (0.27,0.37) |
| 5 | 0.21(0.03) | (0.14,0.26) | 0.27(0.03) | (0.21,0.34) |

**Table S1.** Fast Reaching Experimental data (see *Fast Reaching Task*). For five subjects, we present the mean, standard deviation, minimum and maximum of the movement durations in ten reaches to a large (L) and small (S) square target. The target widths are 8 cm and 2 cm, and the reach amplitudes are 15 cm.

### S1. Materials and Methods

We give the details of the terms $\mathrm{l}(\mathrm{r})$ and $\rho(\mathbf{q}_\mathrm{N})$ in Equation (1). For a joint value r with limits l and u (denoting the lower and upper bounds for the coordinate), $\mathrm{l}(\mathrm{r})$ denotes a quadratic penalty on joint limit violations:

$$\mathrm{l}(\mathrm{r}) = \begin{cases} 0 \ \text{ if } \mathrm{l} < \mathrm{r} < \mathrm{u} \\ (\mathrm{r}-\mathrm{l})^2 \ \text{ if } \mathrm{r} < \mathrm{l} \\ (\mathrm{r}-\mathrm{u})^2 \text{ if } \mathrm{r} > \mathrm{u} \end{cases} \qquad \mathrm{S}(1)$$

The $\rho(\mathbf{q}_\mathrm{N})$ is a quadratic term to encourage forearm pronation at the end of the movement:

$$\rho(\mathbf{q}_\mathrm{N}) = (r_{forearm} - \phi)^2 \qquad \mathrm{S}(2)$$

where $r_{forearm}$ denotes the coordinate of the forearm and $\phi$ is the forearm value denoting pronation ($\pi$ rad in our modeling platform).
